# FUNCTIONAL FRAGMENTATION AND STRUCTURAL DRIVERS OF THALAMO-CORTICAL CIRCUITS IN TEMPORAL LOBE EPILEPSY

**DOI:** 10.64898/2026.07.07.737072

**Authors:** Rui Ding, Ke Xie, Judy Chen, Alexander Ngo, Fatemeh Fadaie, Guan Zhou, Ella Sahlas, Jordan DeKraker, Jessica Royer, Raul Rodriguez-Cruces, Thaera Arafat, Yejin Ann, Seok-Jun Hong, Alexandra John, Sofie Valk, Zhiqiang Zhang, Luis Concha, Paule-J Toussaint, Raluca Pana, Neda Bernasconi, Andrea Bernasconi, Alan C Evans, Boris C. Bernhardt

**Affiliations:** Montreal Neurological Institute, McGill University, Montreal, QC, Canada; Department of Medical Imaging, Jinling Hospital, Nanjing University School of Medicine, Nanjing, China; Institute of Neurobiology, Universidad Nacional Autónoma de Mexico (UNAM), Queretaro, Mexico; Sungkyunkwan University, S.Korea; Max Planck Institute for Human Cognitive and Brain Sciences, Leipzig, Germany

**Author notes:** joint senior authors (Alan C Evans, Boris C. Bernhardt). **Corresponding authors:** Rui Ding and Boris Bernhardt.

**Keywords:** Epilepsy, TLE, connectome, resting-state fMRI, network pathology

## Abstract

**Objective:** In temporal lobe epilepsy (TLE), the thalamus acts as a nexus in a pathophysiological network that implicates mesiotemporal, subcortical, and neocortical regions. Studying a large multimodal and multicentre dataset, we profiled thalamic, hippocampal, and neocortical functional connectivity (FC), assessed structural mediators, and examined clinical associations.

**Methods:** We studied resting-state FC alongside structural and diffusion MRI data in 250 unilateral TLE patients and 259 healthy controls, with measures aggregated across four independent datasets. Data were processed using open-access neuroinformatics workflows and analyzed at a subregional level to maximize anatomical precision. Statistical analysis and mediation models assessed between-group FC changes, structural contributors, and clinical correlations.

**Results:** Compared to controls, TLE patients presented with reduced thalamo-cortical FC, which was most marked in mesiotemporal, fronto-central, and occipital regions. Thalamo-hippocampal FC was also reduced, with effects seen in all CA subfields. In the thalamus, FC reductions peaked in the ventral posterior nucleus when considering neocortical target regions and in the mediodorsal nucleus when considering hippocampal target regions. While ipsilateral hippocampal volume and diffusion changes mediated thalamo-hippocampal FC, thalamo-cortical FC appeared decoupled from structural alterations. Findings were consistent in left and right TLE patients, in patients with short and long disease duration, and across imaging sites, suggesting that thalamo-cortical FC imbalances are a consistent signature of TLE. Conversely, thalamo-hippocampal FC was elevated in patients with focal-to-bilateral-tonic-clonic seizures and FC alterations were more marked in the subgroup of operated patients that became seizure-free after surgery.

**Conclusion:** Our multi-site findings demonstrate marked thalamic circuit fragmentation in TLE. Ipsilateral findings robustly showed subdivision-specific effects, which point to both mesiotemporal co-lateralization as well as broader system-level involvement. Mediation analyses furthermore confirmed a key role of hippocampal pathology in disrupted thalamo-hippocampal connectivity in TLE, while broader thalamo-cortical fragmentation becomes increasingly independent of mesiotemporal compromise. Critically, thalamic FC represents a network substrate for seizure generalization and can serve as a prognostic indicator for surgical outcome. These results underscore the contribution of the thalamus as a hub in macroscale dysfunction in TLE.

## Introduction

The thalamus serves as a central hub that mediates bidirectional communication between cortical and subcortical systems, integrating information across large-scale networks [1]. These relay and modulatory functions are governed by specialized nuclei that maintain distinct connections to both mesiotemporal regions, such as the hippocampus, and broader neocortical networks [2, 3]. Temporal lobe epilepsy (TLE), the most prevalent pharmaco-resistant epilepsy in adults, involves pathological reorganization of these circuits. While TLE is primarily associated with seizures generated in temporo-limbic networks as well as mesiotemporal pathology [4, 5], abundant evidence from histology, neuroimaging, and electrophysiology suggests thalamic involvement [6-8]. Overall, these findings support that the thalamus acts as a critical bridge linking the mesiotemporal epicenter to broader cortical networks.

Neuroimaging and brain connectivity analysis provide powerful *in-vivo* windows into brain organization in both health and disease [9, 10]. In TLE, a robust literature has mapped structural abnormalities, showing relatively consistent evidence for atrophy of the thalamus in patients relative to controls [11, 12]. Several studies have also begun to chart subregional changes in thalamic morphology, highlighting differential alterations across several thalamic divisions [7, 12]. At the level of brain function, while some studies have examined intrinsic functional connectivity (FC) alterations between the thalamus and cortical regions based on resting-state functional MRI (rs-fMRI), few studies have specifically investigated the thalamus as a nexus linking the mesiotemporal disease epicentre with broader neocortical networks in TLE. Furthermore, existing findings show somewhat inconsistent results, with some studies reporting no significant differences, while others reported either decreased or increased FC when comparing patients to controls [13, 14]. These mixed findings likely stem from relatively small samples studied, site-specific variations, differences in acquisition protocols, as well as heterogeneous analytical methodologies. Importantly, prior research has so far largely failed to address in how structural damage relates to functional network disruption though structural alterations play essential roles in shaping FC changes [15].

We conducted a multi-site rs-fMRI study examining alterations in FC between the thalamus and mesiotemporal as well as neocortical networks in TLE patients compared to healthy controls. Firstly, we analyzed the FC between the thalamus and cortical as well as hippocampal subregions, followed by detailed subregional assessment of the thalamus by using the Tian atlas [16], derived from fMRI functional connectivity gradients, this functionally-defined parcellation provides logical consistency with our FC-based analyses at the subregional level, allowing us to identify which thalamic subdivisions and target regions are differentially affected in TLE. To ensure robustness of subregional findings against parcellation choice, we additionally validated key results using the histology-based HIPS-THOMAS atlas [17], which is grounded in anatomically-defined thalamic nuclei. All data were processed through a uniform neuroinformatics workflow with ComBat harmonization to mitigate sites batch effect. Secondly, we assessed whether structural changes, including volumetric as well as microstructure alterations in the thalamus and hippocampus, were related to atypical FC in TLE utilizing mediation analysis, allowing us to test whether structural pathology statistically accounted for FC alterations. Thirdly, leveraging the clinical phenotyping available in our cohort, we evaluated associations with electro-clinical variables including demographics, seizure history, and postoperative outcome in patients who underwent surgery, to assess the clinical relevance of identified FC alterations.

## Methods

### Participants

We analyzed 3T rs-fMRI data from 250 unilateral TLE patients (136 females; mean±SD age=30.85±9.42 years; range=15-58 years; disease duration=13.96±10.75 years) and 259 age- and sex-matched healthy controls (130 females; mean±SD age=29.54±8.42 years; range=17-60 years). Healthy controls and epileptic patients were aggregated across four independent datasets from three different sites: Montreal Neurological Institute and Hospital (*MICA-MICs* [18]; n=118; 46 TLE; *NOEL* n=100; 62 TLE), Jinling Hospital, China (*NKG*; n=226; 113 TLE) [19], Universidad Nacional Autonoma de Mexico (*EpiC*; n=65; 29 TLE) [20].

Patients were diagnosed according to the ILAE classification criteria based on comprehensive evaluation that included clinical history, seizure characteristics, video-EEG recordings, neuropsychological testing, and clinical MRI findings. TLE cohort: At the time of data analysis, 90/250 TLE patients had undergone epilepsy surgery with available post-operative outcome data. MRI data were acquired prior to surgery in all operated patients. Seizure outcome was determined using Engel’s classification with a mean follow-up duration of 54.40±32.01 months (range: 1–120 months). Post-surgically, 65/90 TLE patients (72.2%) achieved seizure freedom (Engel I), while 25 patients (27.8%) experienced ongoing seizures (Engel II-IV). Detailed seizure semiology data were available for 171 patients, including focal to bilateral tonic-clonic seizures (FBTCS; n=83) and focal seizures (FS; n=88). Detailed dataset-specific demographic and clinical information is available in **Table 1**.

**Table 1.** Demographic and clinical information of the datasets.

|  | **MICA-MICS** |  |  | **NKG** |  |  | **EPIC** |  |  | **NOEL** |  |  |
| --- | --- | --- | --- | --- | --- | --- | --- | --- | --- | --- | --- | --- |
|  | *Controls* | *TLE* | *p* | *Controls* | *TLE* | *p* | *Controls* | *TLE* | *p* | *Controls* | *TLE* | *p* |
| *Number* | 72 | 46 |  | 113 | 113 |  | 36 | 29 |  | 38 | 62 |  |
| *Age* | 32.30±7.71<br>(23-60) | 34.34±9.68<br>(19-57) | 0.07<sup>a</sup> | 25.61±5.44<br>(20-40) | 27±7.04<br>(16-43) | 0.10<sup>a</sup> | 33.53±11.75<br>(17-57) | 30.90±11.75<br>(17-58) | 0.38<sup>a</sup> | 32.13±8.11<br>(20-53) | 34.55±8.9<br>(19-52) | 0.18<sup>a</sup> |
| *Sex(M/F)* | 39/33 | 23/23 | 0.66<sup>b</sup> | 60/53 | 59/54 | 0.84<sup>b</sup> | 10/26 | 19/20 | 0.77<sup>b</sup> | 20/18 | 23/39 | 0.11<sup>b</sup> |
| *Focus(L/R)* | - | 28/18 |  | - | 56/57 |  | - | 17/12 |  | - | 30/32 |  |
| *Duration* |  | 15.34±11.3<br>(1-45) |  |  | 11.02±8.79<br>(0.1-37) |  |  | 15.52±13.23<br>(2-49) |  |  | 17.55±11.12<br>(1-41) |  |

Age and disease duration are presented as mean (±SD) years. <sup>a</sup>Two-sample t-test. <sup>b</sup> Chi-square test

#### Standard Protocol Approvals, Registrations, and Patient Consents

Local institutional review boards and ethics committees approved each included cohort study. Written informed consent was provided by all participants (or their guardians) according to local requirements and in accordance with the Declaration of Helsinki.

### Image acquisition

Multimodal neuroimaging data were collected using 3T MRI systems, with protocols including structural T1-weighted imaging, diffusion MRI, and resting-state fMRI. The *EpiC* dataset was acquired on a 3T Philips Achieva scanner. Structural T1-weighted MRI were acquired using a 3D spoiled gradient-echo sequence (1mm³ isovoxels, TR/TE=8.1/3.7ms, FA=8°, FOV=256×256mm²); rs-fMRI using a gradient-echo EPI (2×2×3mm³ voxels, TR/TE=2000/30ms, FA=90°, 34 slices, 200 volumes); and diffusion imaging using a single-shell EPI acquisition (2mm³ isovoxels, TR/TE=11860/64.3ms, b=2000s/mm², 60 directions). The *MICA-MICs* dataset was acquired on a 3T Siemens Prisma-Fit. Structural T1-weighted images were acquired using a 2D MPRAGE sequence (0.8mm³ isovoxels, TR/TE/TI=2300/3.14/900ms, FA=9°, 320×320 matrix); rs-fMRI using a multi-band accelerated 2D gradient-echo EPI (3mm³ isovoxel, TR/TE=600/30ms, FA=52°, acceleration factor=6, 48 slices, 700 volumes); and diffusion imaging using a multi-shell 2D EPI acquisition (1.6mm³ isovoxels, TR/TE=3500/64.4ms, b-values=300/700/2000s/mm² with 10/40/90 directions). The *NKG* dataset was acquired on a 3T Siemens Trio scanner. Structural T1-weighted MRI were acquired using a 3D-MPRAGE (0.5×0.5×1mm³ resolution, TR/TE=2300/2.98ms, FA=9°); rs-fMRI using a 2D-EPI acquisition (3.75×3.75×4mm³ voxels, TR/TE=2000/30ms, FA=90°, 30 slices, 255 volumes); and diffusion MRI using a 2D EPI acquisition (0.94×0.94×3mm³ resolution, TR/TE=6100/93ms, b=1000s/mm², 120 directions). The *NOEL* dataset was acquired on a 3T Siemens Trio scanner. Structural T1-weighted MRI were acquired using a 3D-MPRAGE (1mm³ isovoxels, TR/TE=2300/2.98ms, FA=9°); rs-fMRI using a 2D gradient-echo EPI (4mm³ isovoxels, TR/TE=2020/30ms, FA=90°, 34 slices, 150 volumes); and diffusion imaging using a twice-refocused 2D EPI acquisition (2×2×3mm³ resolution, TR/TE=8400/90ms, FA=90°, b=1000s/mm², 64 directions). Notably, participants were included only if they did not show excessive head motion, defined as mean FD within 2 standard deviations of the group mean (cutoff: <0.35 mm), and did not show low signal-to-noise ratio, defined as tSNR within 2 standard deviations of the group mean (cutoff: >19.29) (**Fig. S1**). TLE patients and controls did not differ in terms of age (*t*=1.66, *p*=0.10) and sex (χ2=0.90, *p*=0.34).

### Multimodal processing

Data were preprocessed using micapipe (v0.2.3; https://micapipe.readthedocs.io/) [21], a multimodal MRI processing and data fusion pipeline, which integrates AFNI, FSL, FreeSurfer, ANTs, and Workbench.

For structural MRI processing, T1-weighted images were de-obliqued and reoriented, followed by linear co-registration, intensity non-uniformity correction, and skull stripping. The thalamus, neocortex, and hippocampus were automatically segmented using FSL 6.0, FreeSurfer 6.0.0, and HippUnfold 1.0.025 [22]. Volumetric measurements were obtained using FreeSurfer for the thalamus and HippUnfold 1.0.025 for the hippocampus [22].

Resting-state fMRI preprocessing involved discarding the first five volumes, followed by image reorientation, slice-timing, and motion correction. Nuisance signals were removed using an in-house trained ICA-FIX classifier for *MICA-MICs* data and linear regression for data from other datasets (white matter and Cerebrospinal Fluid, CSF). Preprocessed fMRI timeseries were co-registered to each individual’s cortical surface, initially resampled to the Conte69 32k vertices per hemisphere, followed by downsampling to 5k vertices/hemisphere. The fMRI timeseries were also mapped onto the hippocampal midthickness surface derived from HippUnfold [22]. Volumetric fMRI timeseries were aligned to MNI152 space in 2mm resolution to extract whole-thalamus and individual thalamic division timeseries. The atlas used for the thalamus parcellation was based on the fMRI FC gradient derived from an independent sample and enabled the quantification of subcortical FC gradients to guide boundary delineation [16]. We selected this atlas for three reasons. First, as our primary outcome was thalamic FC, a functionally defined parcellation ensures consistency between the parcellation strategy and the analytic target. Second, the Tian atlas is designed for direct integration with cortical atlases, enabling coherent thalamo-cortical analyses. Third, it offers multi-scale granularity and has demonstrated reliability in 3T multi-site data, matching our study design. The thalamus was divided into 8 subdivisions: Lateral dorsoanterior thalamus (THA-DAl); Medial dorsoanterior thalamus (THA-DAm); Superior ventroanterior thalamus (THA-VAs); Inferior ventroanterior thalamus:Anterior division (THA-VAia), Posterior division (THA-VAip); Dorsoposterior thalamus (THA-DP); Medial ventroposterior thalamus (THA-VPm); Lateral ventroposterior thalamus (THA-VPl). This atlas demonstrates high spatial correspondence with histological atlases, with specific subdivisions aligning with established anatomical nuclei: VAip corresponds to the mediodorsal nucleus, VAia to the ventral anterior nucleus, VPm to the parafascicular nucleus, VPl to the ventral posterior lateral nucleus, VAs to the ventral anterior nucleus, DAm to the lateral posterior nucleus, DAl to the central lateral nucleus, and DP to the anterior pulvinar. Thalamic segmentation was additionally performed using the HIPS-THOMAS pipeline, a histologically-defined multi-atlas segmentation method [17]. Native T1-weighted images were first synthesized into WMn-MPRAGE-like contrast using the HIPS-THOMAS Docker implementation (https://github.com/thalamicseg/hipsthomasdocker) to enhance thalamic nuclei delineation. The fMRI timeseries were then coregistered from native functional space into T1 space, and timeseries were extracted from ten THOMAS nuclei: anteroventral (AV), ventral anterior (VA), ventral lateral anterior (VLa), ventral lateral posterior (VLp), ventral posterior lateral (VPL), pulvinar (Pul), lateral geniculate (LGN), medial geniculate (MGN), centromedian (CM), and mediodorsal (MD).Thalamic timeseries were extracted at two levels: global thalamic timeseries were computed by averaging fMRI signals across all thalamic voxels, while subdivision-specific timeseries were calculated by separately averaging fMRI signals within each of the eight/ten thalamic subdivisions. Spatial smoothing was applied with kernels of 10mm full width at half maximum (FWHM) for cortical surfaces, 3mm FWHM for hippocampus). MRI quality control procedures included evaluating signal-to-noise ratio and visual inspection of surface extractions for structural MRI, as well as analyzing temporal derivatives and framewise displacement metrics for resting-state fMRI (see: **Supplementary Materials**).

### FC analysis

FC was calculated as the product moment correlation between thalamic fMRI timeseries (both global and subdivision-specific) and vertex-wise timeseries from the neocortex and hippocampus (**Fig. 1A**). Fisher’s r-to-z transformations normalized FC distributions. To mitigate multi-site batch effects, we employed ComBat harmonization [23], which corrects for site-specific variance while preserving biologically relevant variance. Subsequently, left and right hemisphere of TLE patients were sorted relative to the epileptogenic focus (*i.e.,* into ipsilateral and contralateral). To account for normal inter-hemispheric asymmetries, prior to sorting we normalized FC using z-transformations relative to healthy controls [24]. Statistical comparisons of FC patterns between TLE patients and controls were performed using linear models implemented in BrainStat (http://brainstat.readthedocs.io) [25], with age and sex included as covariates. Analyses were conducted at both global thalamic and thalamic subdivision levels. Effect sizes were quantified using Cohen’s *d*. Multiple comparison were corrected using random field theory (RFT) with family-wise error rate, FWER [26] control for cortical regions and permutation with FWER correction for hippocampal regions.

**Fig. 1.**
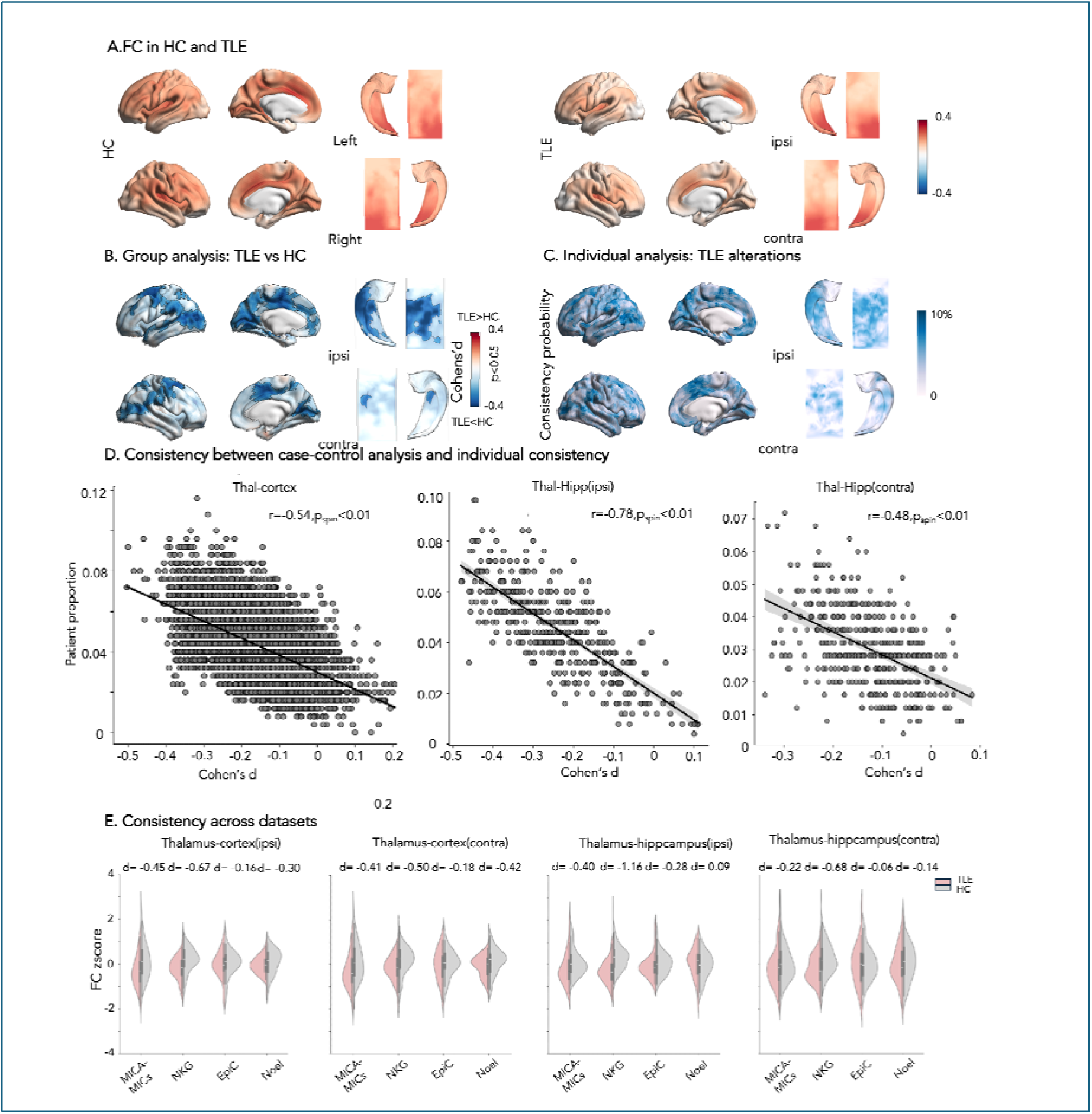
Between-group differences in functional connectivity (FC) between global thalamus and cortex/hippocampus. **(A)** FC patterns in HC (n=259) and TLE patients (n=250). **(B)** Generalized linear models, controlling for age and sex revealed significant FC differences between groups, after data have been processed using equivalent workflows and harmonized across sites. Results were corrected for multiple comparisons using random field theory correction for cortical regions and permutation test for hippocampal regions, controlling for the family-wise error rate (*p*_FWER_<0.05). Significant clusters are outlined in black. **(C)** Individual consistency was assessed by calculating the percentage of patients whose z-transformed FC values (relative to HC) fell below *z<-1.96* at each vertex, classified as abnormal. Results showed similar patterns to the group-level case-control analysis. **(D)** Consistency between case-control analysis (Cohen’s d) and individual consistency (proportion of patients with z-scores <-1.96). **(E)** Cross-dataset consistency analysis. Mean FC values were calculated for each dataset within regions showing significant alterations, and TLE vs. HC differences were compared across datasets. ***Abbreviations***: *HC*, healthy controls; *TLE*, temporal lobe epilepsy; *ipsi*, ipsilateral; *contra*, contralateral.

*Post-hoc* analyses examined the consistency of findings across datasets. We evaluated inter-dataset consistency by extracting vertices that showed significant FC differences in the main multi-site analysis and computing their mean connectivity values in each dataset. We then charted effect sizes for TLE *vs* control differences across each of the four datasets individually. We additionally examined consistency at the individual patient level [27]. Each TLE patient’s FC values were z-transformed relative to the distribution in healthy controls. We then quantified the proportion of patients exceeding *z<-1.96* within a region. This resulted in maps indicating the percentage of patients showing extreme FC reductions. Then, we used spin permutation [28] to test the consistency between case-control analysis and individual consistency results.

For analysis of thalamic subdivisions, we employed a parallel approach to the global thalamic analysis. First, FC was computed between each of the eight thalamic subdivisions and cortical/hippocampal regions. We identified cortical and hippocampal regions showing significant FC alterations across subdivisions and generated maps indicating the number of subdivisions demonstrating significant FC changes within each region. For each thalamic subdivision, we calculated the mean FC values within regions identified as significant in the multi-site analysis. FDR correction [29] were applied across the eight thalamic subdivisions. Individual patient consistency was assessed following the same procedure as the global thalamic FC analysis, applying a threshold of *z<-1.96*. The proportion of patients exceeding this threshold was calculated separately for ipsilateral and contralateral cortical and hippocampal regions within the significant clusters identified in the multi-site analysis.

Furthermore, we conducted targeted analyses to establish more precise functional correspondence between thalamic subdivisions and specific brain systems. Mean timeseries were extracted from individual hippocampal subfields and Yeo 7-network parcellations [30], which were then correlated with each thalamic subdivision timeseries using product moment correlations following identical statistical procedures as the multi-site analysis. FDR correction [29] controlled for multiple comparisons across these targeted FC analyses.

### Assessment of structural alterations

To assess the role of structural changes in FC changes, we conducted mediation analyses. Global thalamic and hippocampal volumes were first adjusted for estimated total intracranial volume. Subsequently, both volume and mean diffusivity (MD) metrics were harmonized using ComBat [23] to mitigate multi-site batch effects. Finally, all metrics were z-normalized relative to healthy controls and categorized into ipsilateral and contralateral hemispheres [24]. We focused on mean diffusivity (MD) as a comprehensive index of microstructure, which has been frequently observed to be increased in TLE patients in prior work [31]. In the mediation models, group (TLE *vs* HC) served as the independent variable, structural measures as potential mediators (bilateral thalamic volumes, ipsilateral hippocampal volume, bilateral thalamic mean diffusivity (MD), and ipsilateral hippocampal MD), and mean FC within significantly altered regions in multisite analysis between thalamus and cortex/hippocampus as the dependent variable. The significance of indirect effects was assessed using bias-corrected bootstrap confidence intervals with 1000 iterations. To account for multiple testing across these mediation models, p-values were corrected using FDR adjustment.

### Clinical association analysis

Several analyses examined associations to clinical variables in TLE.

*a) Lateralization effects.* We separately analyzed left (n = 131) and right TLE (n = 119) to explore seizure focus lateralization effects. Vertex-wise analyses were performed on ComBat harmonized data, controlled for age and sex.
*b) Correlation to electro-clinical variables and post-operative seizure outcome.* To explore progressive FC changes, we examined disease duration correlations with FC across harmonized data (n=250) from *MICA-MICs*, *NKG,* and *NOEL* datasets, controlling for sex. We also examined differences in harmonized FC across clinical subgroups, comparing focal to bilateral tonic-clonic seizures (FBTCS, n = 83) and focal seizures (FS, n = 88) in the *MICA-MICs, NKG* and *NOEL* datasets. In the subgroup of operated patients from the *MICA-MICs, NKG*, and *NOEL* datasets, we also compared the patients that will become seizure-free (SF, n = 65) *vs* non-seizure-free (NSF, n = 25) after their surgery. Given that hippocampal sclerosis patients and shorter duration patients typically show favorable surgical outcomes [32, 33], we assessed whether the FC trend was confounded by hippocampal volume and disease duration using ANCOVA.

## RESULTS

### Broad functional fragmentation of thalamic circuits in TLE

In healthy controls, the FC between the thalamus and cortex exhibited a relatively symmetric pattern, with broadly high connectivity values throughout the cortex that were highest in prefrontal and parietal regions. Similarly, FC between the thalamus and hippocampus demonstrated a symmetric pattern of connections across hemispheres that was highest in posterior regions (**Fig. 1A**).

Comparing TLE patients to controls, we observed marked reductions in global thalamo-cortical and thalamo-hippocampal FC, indicative of a broad fragmentation of functional circuits. Neocortical FC reductions were primarily observed in the ipsilateral prefrontal, precentral, postcentral, parietal, visual cortices, and medial temporal regions, while hippocampal FC reductions were most prominent across ipsilateral CA subfields (Cortex: ipsilateral: *p*_FWER_<0.05, mean±SD Cohen’s *d* in significant clusters: - 0.27±0.05; contralateral: -0.26±0.04; hippocampus: ipsilateral: *p*_FWER_<0.05, -0.33±0.07; contralateral:-0.28±0.04, **Fig. 1B**). Individual consistency analysis calculated the proportion of TLE patients showing FC alterations in each region compared to controls (**Fig. 1C**). The individualized results were highly congruent with the group-level case-control analysis, with higher proportions of patients showing alterations in prefrontal, precentral, postcentral, parietal, and visual cortices as well as in hippocampal subfields. The spin test results also showed consistent pattern between group analysis and individual analysis (**Fig. 1D**). Despite variability in effect sizes, the overall direction of FC reductions was consistent across all datasets (thalamus-cortex: ipsilateral: *MICA-MICs/NKG/EpiC/NOEL d*=−0.45/−0.67/−0.16/-0.3; contralateral: *d*=-0.41/−0.50/−0.18/-0.42;thalamus-hippocampus:ipsilateral: *d*=−0.40/−1.16/−0.28/0.09; contralateral: *d*=0.22/−0.68/−0.06/-0.14,**Fig. 1E**).

Analysis of thalamic subnuclei revealed an ipsilateral dominated reduction in FC to neocortical systems as well as the hippocampus (**Fig. 2B**). Effects in contralateral divisions were more subtle. Considering thalamo-cortical connections, ventroposterior subdivisions (VPL, VPM, VAip) showed the most marked FC reductions to ipsilateral neocortical systems (**Fig. 2C**). Considering thalamo-hippocampal connections, all subdivisions were affected but VAip exhibited the greatest FC decrease to ipsilateral hippocampal subregions (**Fig. 2C**). Individual patient analyses showed patterns consistent with group-level case-control findings (**Fig. 2D**). Subdivision-specific brain system analyses revealed distinct patterns of FC disruption. For hippocampal connections, while the disruptions were extended across multiple thalamo-hippocampal pathways, the ipsilateral VAip-CA1 (**Fig. 2E**) pathway showed the most substantial FC decrease compared to other thalamo-hippocampal connections. Thalamo-cortical FC reductions had broader, system-level involvement. VPL-dorsal attention network (DAN) connectivity showed the most pronounced reduction, within a generalized pattern of decreased FC across VP and multiple cortical networks (**Fig. 2F**). We replicated key subregional findings using the histology-based HIPS-THOMAS atlas (**Fig. S2.A**). Consistent with our primary analyses, the mediodorsal nucleus, which has high spatial correspondence to the Tian VAip subdivision, showed the most pronounced FC reductions to ipsilateral hippocampal subregions, with peak alteration at MD-CA1 (**Fig. S2.E**). Thalamo-cortical FC reduction were broadly observed across multiple THOMAS nuclei (notably VLP, VPL, VLa). This wider statistical detection likely reflects the finer nuclei level granularity of HIPS-THOMAS relative to the eight-subdivision Tian atlas. Despite this difference, both parcellations consistently identified the dorsal attention network as a primary site of thalamo-cortical disruption, with the most pronounced reductions involving ventroposterior thalamic regions (**Fig. S2.F**).

**Fig. 2.**
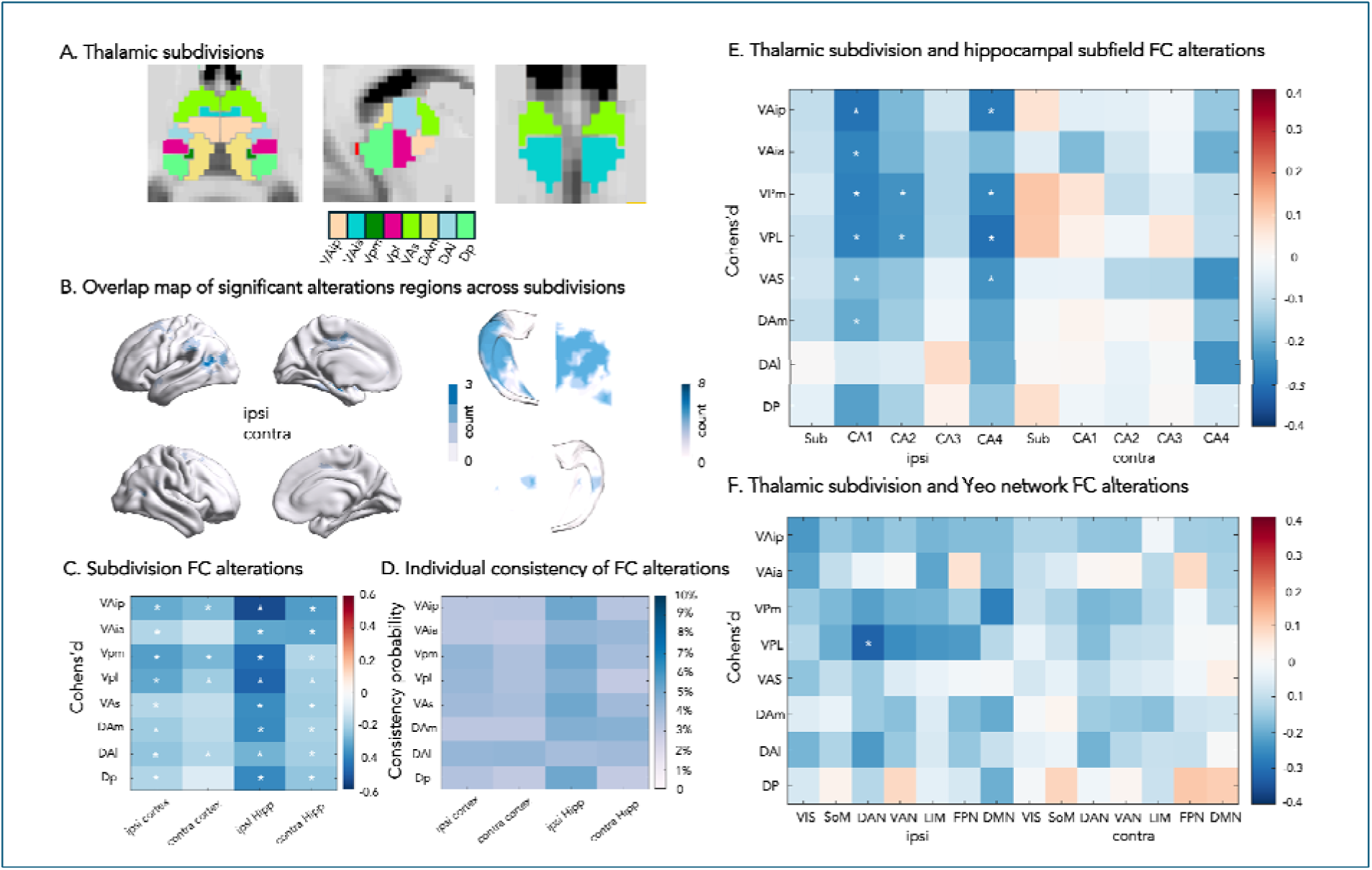
Between-group differences in FC. **(A)** The atlas used in this study divides the thalamus into eight functional subdivisions [16]. **(B)** Overlap maps of significant thalamo-cortical FC differences (TLE *vs* HC) across eight thalamic subdivisions. Color intensity indicates the number of subdivisions showing significant effects. (**C)** Generalized linear models, controlling for age and sex, showed FC differences between two groups across the eight thalamic subdivisions, by calculating the mean value among significant regions in the main analysis. Ipsilateral VPL/VPM and VAip (corresponds to mediodorsal nucleus) showed most pronounced reductions in cortical and hippocampal FC. **(D)** Individual-level consistency, in terms of % of patients with *z<-1.96* relative to controls in each vertex. Mean values were calculated for significant regions from the main analysis, showing similar patterns t the group-level analysis (Fig. 2C). **(E)** Thalamic subdivision-hippocampal subfield FC alterations in TLE, demonstrating the greatest reduction between ipsilateral VAip and CA. **(F)** Thalamic subdivision-Yeo network FC alterations in TLE, with VPL and VPM showing distributed reductions across DAN and DMN. Asterisks denote significance after FDR correction (\**p*_FDR_<0.05) ***Abbreviations***: *VAip*: Inferior ventroanterior-posterior; *VAia* Inferior ventroanterior-anterior; *VPm*: Medial ventroposterior; *VPL*: Lateral ventroposterior; *VAs*: Superior ventroanterior; *Dam*: Medial dorsoanterior; *DAL*: Lateral dorsoanterior; *DP*: Dorsoposterior.

### Associations to structural compromise

TLE patients showed bilateral, yet more marked ipsilateral, thalamic volume reductions (ipsilateral: *d*=-0.64, *p*<0.01; contralateral: *d*=-0.3, *p*<0.01; **Fig. 3A**) and a similar pattern of MD increases (ipsilateral: *d*=0.43, *p*<0.01; contralateral: *d*=0.29, *p*<0.01). Hippocampal volume was reduced and MD was increased ipsilaterally (volume: *d*=-1.29, *p*<0.01; MD: *d*=1.21, *p*<0.01) but not contralaterally (*d*=-0.04, *p*=0.65; *d*=0.17, *p*=0.06).

**Fig. 3.**
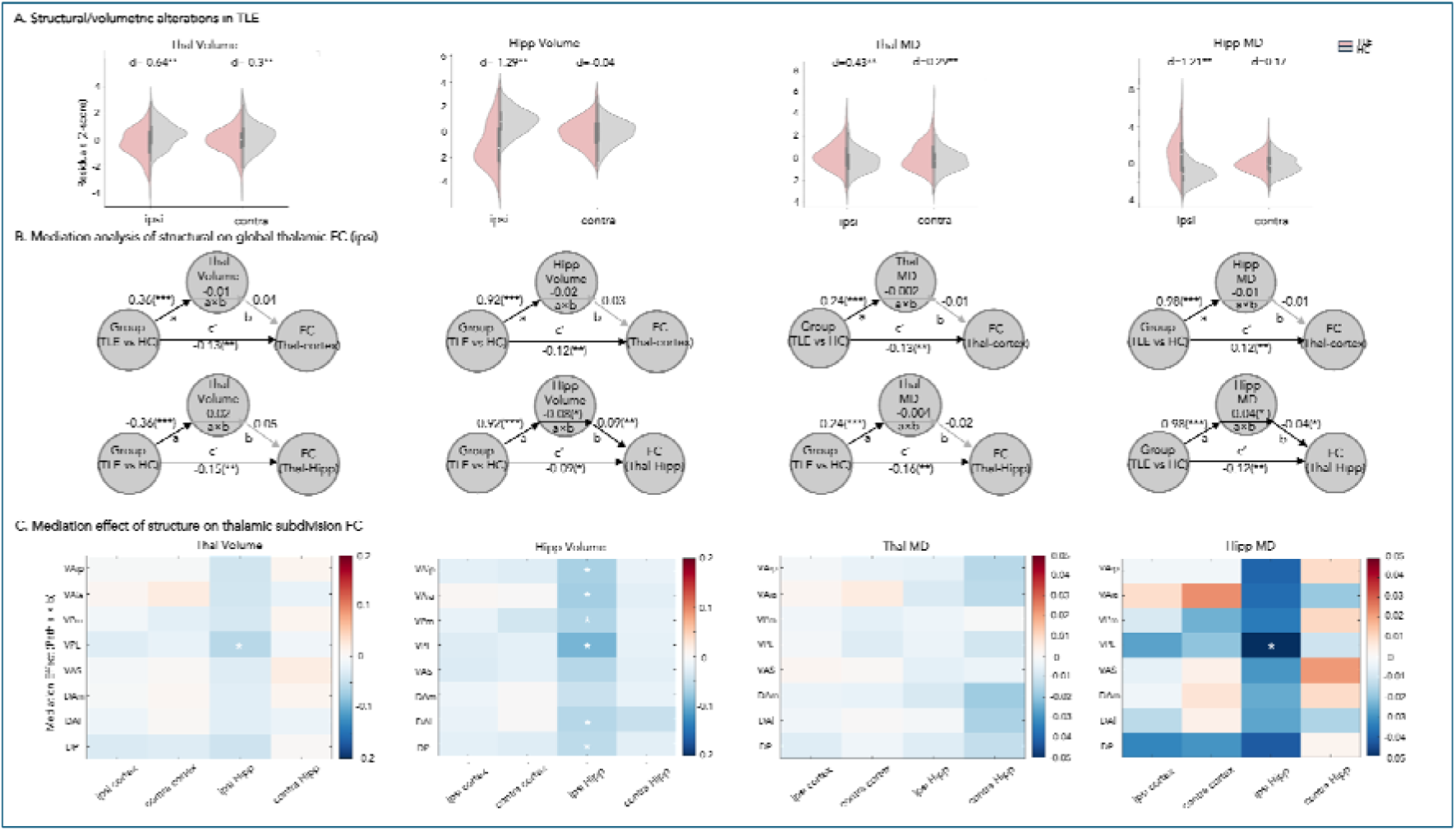
Structure-function relationship. **(A)** Volumetric alterations in thalamus and hippocampus in TLE patients compared to HC. **(B)** Mediation analyses using *group* (TLE *v*s HC) as the predictor, *structure* (thalamic/hippocampal volume, MD) as the mediator, and mean ipsilateral *FC* in significantly altered regions as the dependent variable. Path *a* indicated the effects of *group* on *structure*, path *b* the effects of *structure* on *FC*, and paths *c*’ and *a*×*b* the direct and the indirect/mediation effects of *group* on FC, respectively. *** *p*<0.0005, ** *p*<0.005, * *p*<0.05. **(C)** Mediation effect of the structural changes on subdivision FC alterations. ***Abbreviations***: *HC*, healthy controls; *TLE*, temporal lobe epilepsy; *Thal:* Thalamus. *Hipp:* hippocampus. *ipsi*, ipsilateral; *contra*, contralateral.

Several statistical mediation analyses revealed distinct patterns for thalamic and hippocampal structural contributions to FC disruptions. Considering thalamo-cortical FC patterns, no structural metric significantly mediated these connections (all |mediation effect|<0.02, *p_FDR_*>0.42). In contrast, thalamo-hippocampal FC was partially mediated by hippocampal integrity. Specifically, ipsilateral hippocampal volume showed a significant partial mediation in ipsilateral FC (mediation effect=-0.08, *p_FDR_*<0.05). Additionally, ipsilateral hippocampal MD also demonstrated partial mediation (mediation effect=-0.04*, p_FDR_*<0.005). Notably, neither thalamic volume nor thalamic MD significantly mediated FC difference (all |mediation effects|<0.02, *p_FDR_*>0.2) (**Fig. 3B; contralateral: Fig. S3**).

Analyses at the subdivision level largely replicated the global findings: No structural metrics mediated FC between thalamo-cortical subdivision FC, whereas hippocampal volume significantly mediated thalamo-hippocampal subdivision FC (**Fig. 3C; Thomas: Fig. S4**).

### Clinical associations

Several analyses suggested that thalamic functional fragmentation was consistently observed across the TLE spectrum, with however notable associations to secondary seizure generalization and post-surgical prognosis. Firstly, both left and right TLE patients showed similarly reduced thalamo-cortical and thalamo-hippocampal FC compared to controls, with more pronounced reductions ipsilaterally (Cohen’s *d* range: -0.27 to -0.47; *p*<0.05) (**Fig. S5**). Secondly, we did not observe any significant correlations between duration of epilepsy and thalamic FC (strongest correlation: contralateral thalamo-cortex, *r*=-0.05, *p*=0.40) (**Fig. S6**), suggesting that thalamic FC changes remain consistent across the disease course in patients. On the other hand, patients with a history of FBTCs showed higher thalamo-hippocampal FC (ipsilateral: *d*=0.32, *p*=0.04; contralateral: *d*=0.34, *p*=0.03), with no significant differences in thalamo-cortical (all *d*<0.17, *p*>0.27) (**Fig. S7A**). Ipsilateral thalamo-cortical FC showed a trend toward lower FC in patients that became seizure free after surgery compared to non-seizure free ones (*d*=-0.42, *p*=0.08), with no differences in other connections (all |*d*|<0.17, *p*>0.46) (**Fig. S7B i**). While the latter trend weakened, it persisted after controlling for disease duration and hippocampal volume (*d*=-0.38, *p*=0.11) **(Fig. S7B ii)**.

## DISCUSSION

Leveraging a multimodal high-resolution MRI profiling approach in a large multi-site cohort of TLE patients and healthy controls, we discovered marked evidence for fragmentation of thalamo-cortical and thalamo-hippocampal circuits. FC reductions were particularly marked between the thalamus and ipsilateral prefrontal, sensorimotor, mesiotemporal and visual neocortices as well as ipsilateral CA subfields. Findings were consistent across imaging sites and at the individual patient level. All thalamic nuclei showed predominantly ipsilateral reductions in both cortical and hippocampal FC. However, nucleus-specific patterns emerged, with ventroposterior subdivisions showing greater fragmentation in connectivity to cortical regions compared to other nuclei, while VAip showed the most pronounced hippocampal FC decreases, particularly in VAip-CA pathways. Integration of structural and diffusion MRI measures revealed that thalamo-hippocampal FC alterations were mediated by hippocampal atrophy and microstructural alterations. Findings were similar in left and right TLE patients and consistent with respect to disease duration. On the other hand, we observed higher thalamo-hippocampal FC in patients with a history of FBTCS compared with patients without reported seizure generalization. In addition, we identified a trend towards reduced thalamo-cortical FC in patients that will become seizure free after surgery relative to patients that will experience a post-op seizure relapse. Taken together, our study highlights large-scale hippocampal-thalamo-cortical FC disruptions in TLE that are partially mediated by hippocampal structural compromise. Moreover, our findings suggest that thalamo-hippocampal and thalamo-cortical functional network integrity collectively index patient-to-patient risk in terms of seizure generalization and post-operative seizure recurrence, respectively.

To mitigate potential inconsistencies in thalamo-cortical connectivity patterns across cohorts [13, 14], the current work leveraged a large multisite dataset that was processed using unified neuroinformatics workflows based on open access tools [21, 22] and harmonized using ComBat to remove site-related variance with subregional precision at the thalamic, neocortical, and hippocampal level. Our analysis revealed both global and subdivision-specific patterns of FC disruption in TLE patients and controls. At the global level, thalamo-cortical FC was reduced to frontal-central, parietal, and mesiotemporal target regions compared to healthy controls. With respect to the mesiotemporal lobe, thalamo-hippocampal FC declined predominantly towards CA subfields, hippocampal subregions commonly affected in TLE [4]. Patterns were of variable effect size across datasets, yet in a consistent direction. Findings were also evident in most individual patients, confirming that network disruptions consistently extend beyond the mesiotemporal epicenter. The observed FC alterations are compatible with multiple pathophysiological mechanisms, including disruption of intrinsic network function [34], hypoperfusion and hypometabolism [35], as well as white matter tract disruption [36]. At the subdivision-specific level, ipsilateral VPL and VPM demonstrated relatively greater cortical FC reductions, affecting dorsal attention and default mode systems, while the posterior ventroanterior thalamus nucleus (VAip, which has high spatial correspondence to the mediodorsal nucleus [16]), showed the most pronounced hippocampal FC decline, primarily involving the CA1 subfield but also extending to CA2 and CA4. Validation analyses using the histology-based HIPS-THOMAS atlas confirmed this pattern, identifying the mediodorsal nucleus as the most affected nucleus with peak FC reduction at MD-CA1.These patterns reflect the functional architecture of differently projecting thalamocortical subsystems [37]. VPL and VPM, the ventroposterior nuclei serving as primary somatosensory relays to primary somatosensory cortex [37], showed connectivity disruptions extending beyond their anatomical targets to broader cortical networks, potentially reflecting cascading disruption along interconnected thalamo-cortical systems [38]. The mediodorsal nucleus, projecting densely to hippocampal CA1 [39, 40], showed FC decline that aligned with diffusion MRI evidence of disrupted mediodorsal-hippocampal structural connectivity in TLE [36]. Collectively, these findings establish subdivision-specific thalamic network alterations as a reproducible feature of TLE, robust not only across MRI acquisition parameters and patient populations, but also across functionally- (Tian) and histologically-defined (HIPS-THOMAS) thalamic parcellation schemes.

Considering distributed structural compromise in TLE, it is critical to understand structural contributions to functional network fragmentation. Previous histological reports have shown that TLE pathology extends beyond hippocampus to include cell loss and gliosis of thalamus [41], and neuroimaging investigations revealed both volumetric and microstructural alterations [31, 42]. We investigated whether thalamic and hippocampal volume loss or microstructural alterations mediated the observed FC reductions, revealing differential patterns across network types. Our results confirmed decreased thalamic and ipsilateral hippocampal volumes, as well as increased MD in both structures in TLE, consistent with prior work [31, 42]. Mediation analyses revealed that ipsilateral hippocampal pathology, here indexed via both measures, contribute to thalamo-hippocampal FC disruption. This suggested that both macrostructural atrophy and microstructural tissue alterations (such as cell loss, gliosis, and expanded extracellular spacing [43]) within the epileptogenic hippocampus act as primary drivers of circuit fragmentation between the mesiotemporal lobe and the thalamus. In contrast, neither thalamic nor hippocampal structural measures mediated thalamo-cortical FC, suggesting that these FC disruptions may reflect broader mechanisms beyond regional gray matter damage, potentially involving alterations in white matter tracts [44]. Together, these patterns characterize the thalamus as a broader hub with heterogeneous, circuit-specific pathology. In turn, hippocampal damage likely drives limbic circuit fragmentation, while thalamo-cortical disruptions appear decoupled from regional grey matter structure, likely reflecting broader network reorganization.

We explored the clinical relevance of the FC findings by studying associations with seizure focus laterality, disease duration, history of FBTCs, and post-surgical seizure outcome. Both left and right TLE showed similarly reduced thalamo-cortical and thalamo-hippocampal FC compared to controls, suggesting a common network fragmentation irrespective of the side of seizure onset. Similarly, the absence of a marked duration effect may support that thalamic functional network disruptions remain relatively stable across the disease course [45]. Future longitudinal studies, ideally including patients with new onset of TLE as well as those with at chronic disease stages, are needed to clarify whether findings are already seen early on, and how they progress with time at the within-patient level [45]. Notably, patients with a history of FBTCS showed higher thalamo-hippocampal FC, suggesting that thalamus facilitates bilateral seizure propagation through enhanced hippocampal connectivity, supported by the evidence showed increased thalamo-temporal FC in FBTCS [46] and therapeutic effects of thalamic stimulation desynchronizing this pathway [47].Considering associations between preoperative MRI measures and post-operative seizure freedom, previous literature has already suggested that some cortical rs-fMRI alterations [48] and thalamic structural alterations [49] relate to surgical outcomes in TLE. Here, we extended this literature by showing that seizure-free patients trended towards having lower ipsilateral thalamo-cortical FC compared to non-seizure-free patients. These findings may be seen as consistent with literature suggesting greater "thalamic hubness" predicting seizure recurrence post-op [50], which potentially reflects more marked network reorganization in patients with better surgical prognosis. Importantly, this association persisted after controlling for disease duration and hippocampal volume, indicating that thalamic network alterations carry prognostic information independent of disease chronicity and structural hippocampal damage, two otherwise common correlates of seizure freedom in prior TLE work [32, 33].

## Funding

R.D is funded by China Scholarship Council. K.X. is funded by the China Scholarship Council and the Healthy Brains and Healthy Lives (HBHL). J.C. and E.S. are funded by the Canadian Institutes of Health Research (CIHR) Vanier Canada Graduate Scholarships. A.N. is funded by the Fonds de Recherche du Québec – Santé (FRQS) and the Canadian Institutes of Health Research (CIHR). F.F. is funded by Savoy Foundation for Epilepsy. J.D. was supported by a Natural Science and Engineering Research Council of Canada Post Doctoral Fellowship award (NSERC-PDF). J.R. is funded by the Canadian Institutes of Health Research (CIHR). R.R.C. is funded by the Fonds de Recherche du Québec—Santé (FRQ-S). G.Z. is funded by China Scholarship Council. T.A is funded by Savoy Foundation Postdoctoral Fellowship. A.C.E. acknowledges research support from the CFREF/HBHL. B.C.B. acknowledges research support from the National Science and Engineering Research Council of Canada (NSERC Discovery-1304413), CIHR (FDN-153198, PJT-174995), SickKids Foundation (NI17-039), Helmholtz International BigBrain Analytics and Learning Laboratory (HIBALL), HBHL, Brain Canada Foundation, FRQS, the Tier-2 Canada Research Chairs Program, and the Centre for Excellence in Epilepsy at the Neuro (CEEN).

## Funding

J.D. and B.C.B. are co-founders of BrainScores and hold stock.

## Author contributions

Study Concept/Design: R.D., A.C.E., B.C.B. Data Acquisition, Analysis, Interpretation: R.D.,K.X., J.C.,E.S., A.N., J.D.,T.A., R.R.C., F.F., Y.J.A, P.J.T and B.C.B. Writing: Original Draft: R.D. and B.C.B. Writing – Review & Editing: K.X., J.C., A.N., E.S., J.R., F.F., G.Z.,S.J.H., A.J., S.V., Z. Z, L.C., P.J.T., R.P., N.B., A.B., A. C.E., and B.C.B. Resources: Z.Z., Lu.C., A.B., N.B., and B.C.B. Supervision: A.C.E. and B.C.B.

---

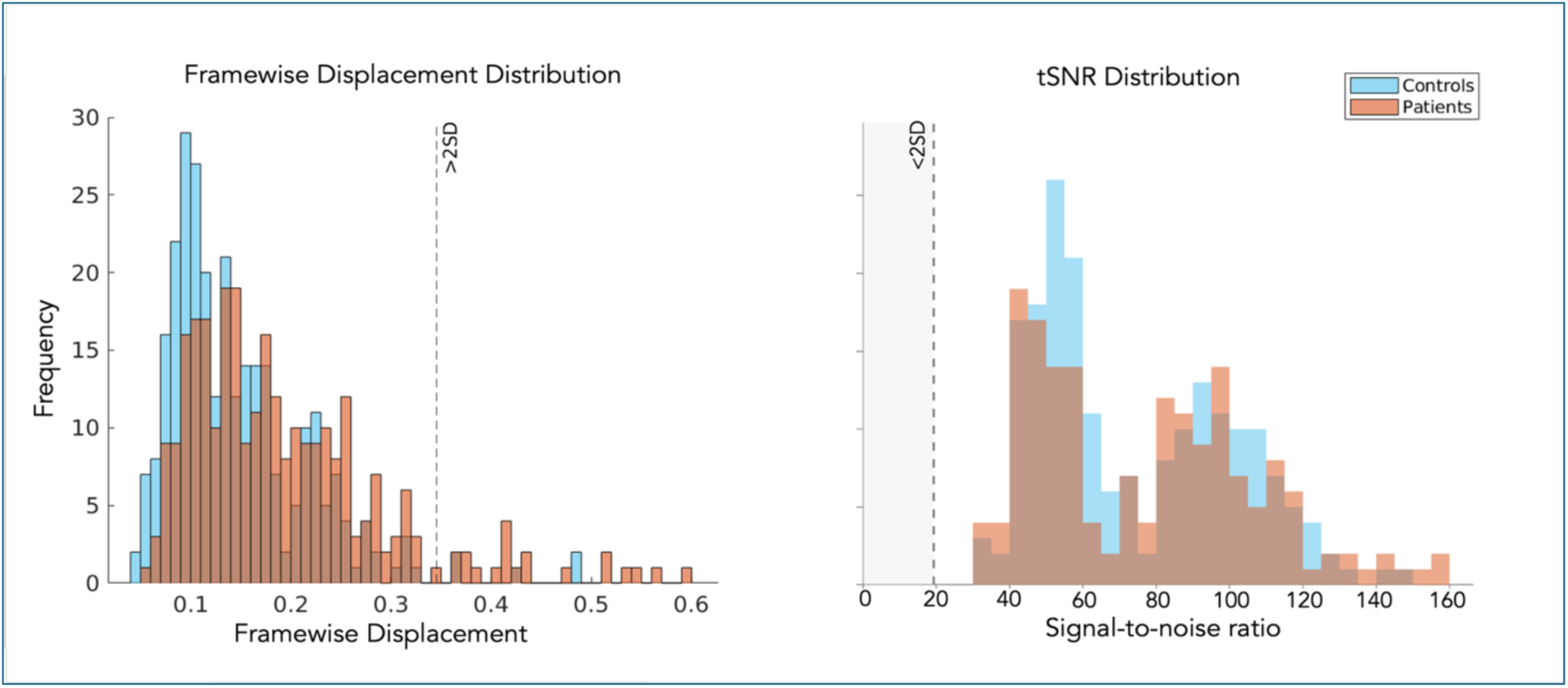

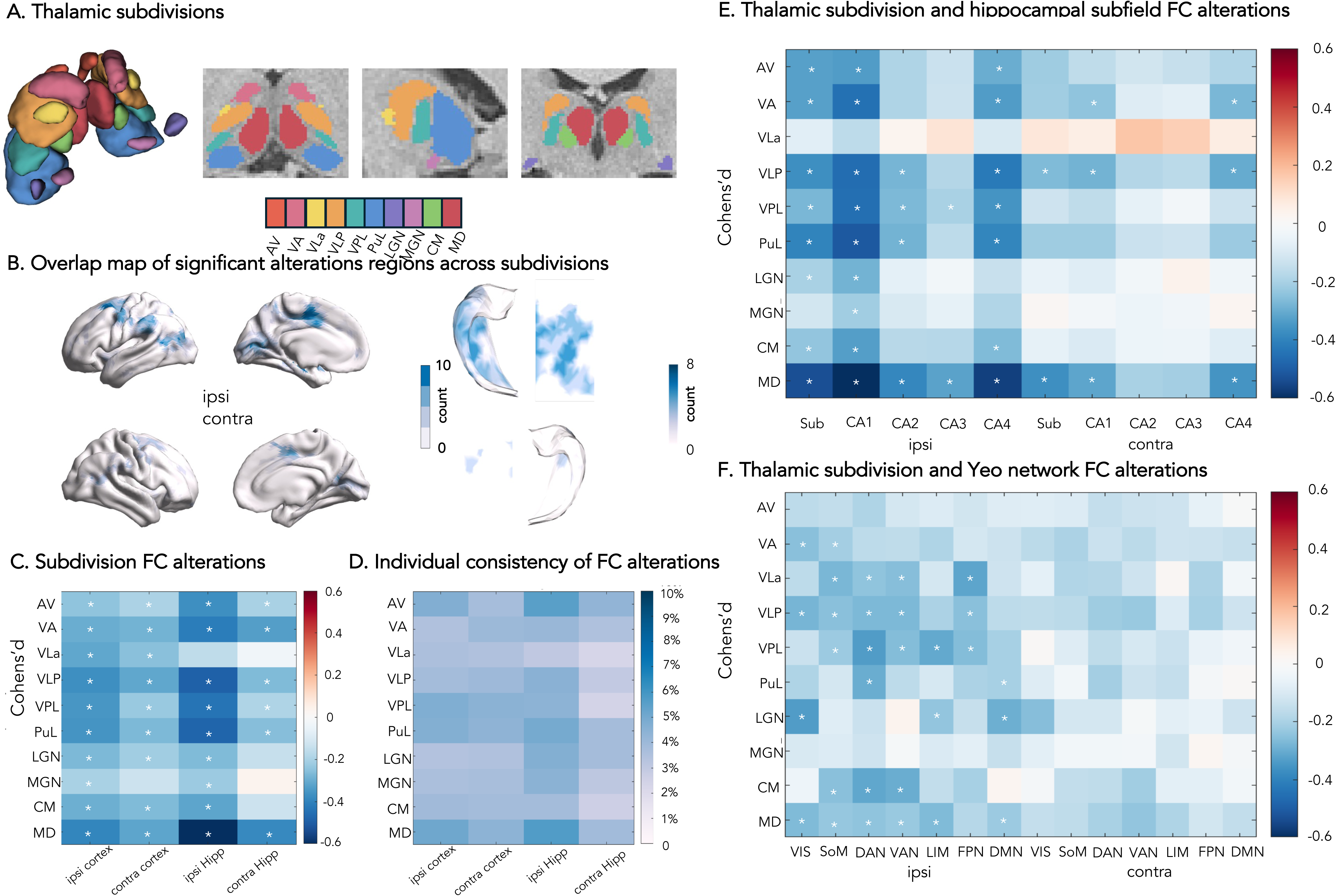

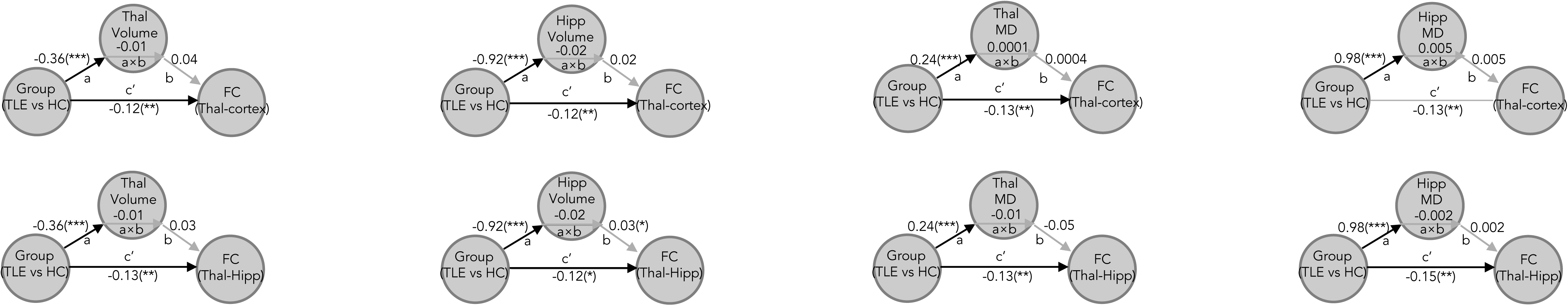

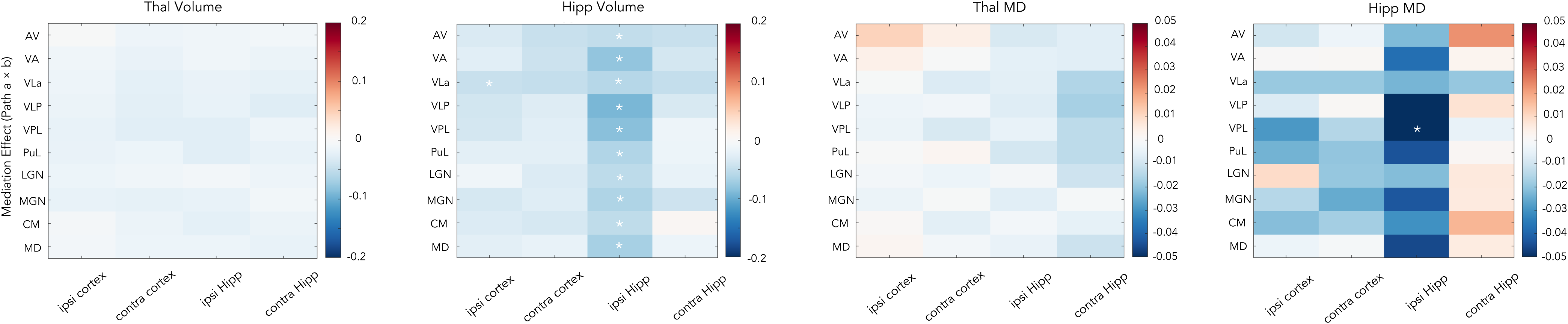

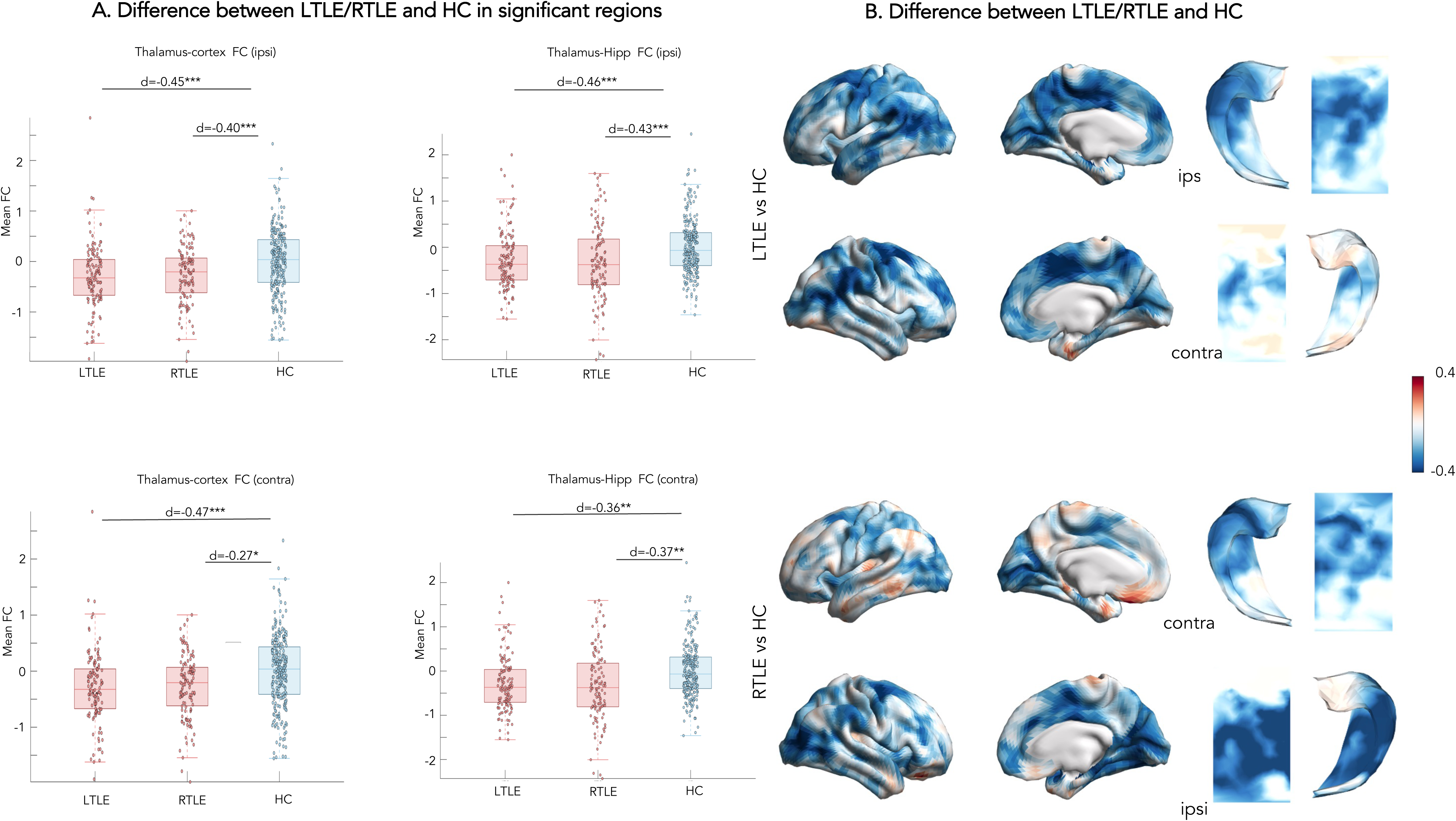

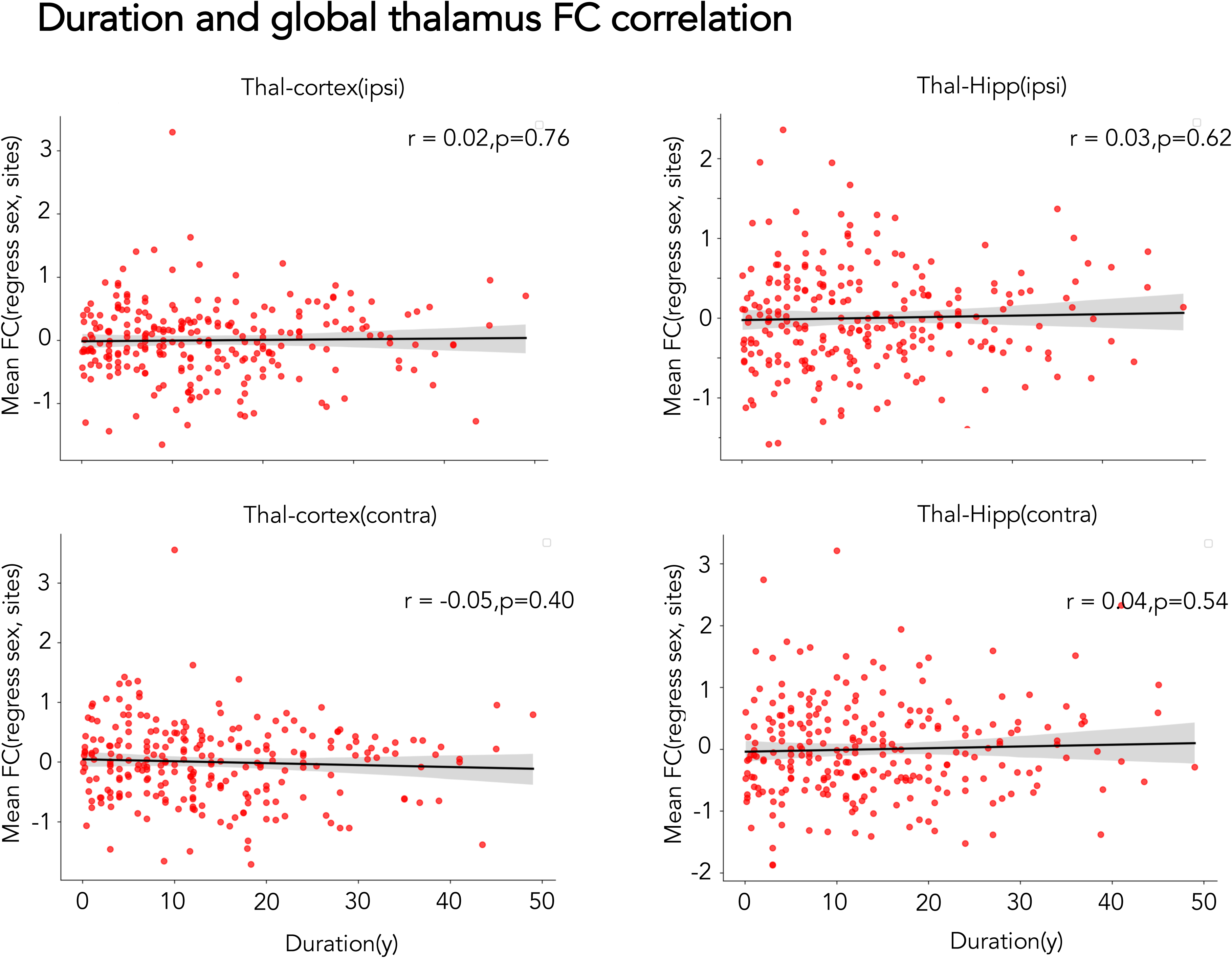

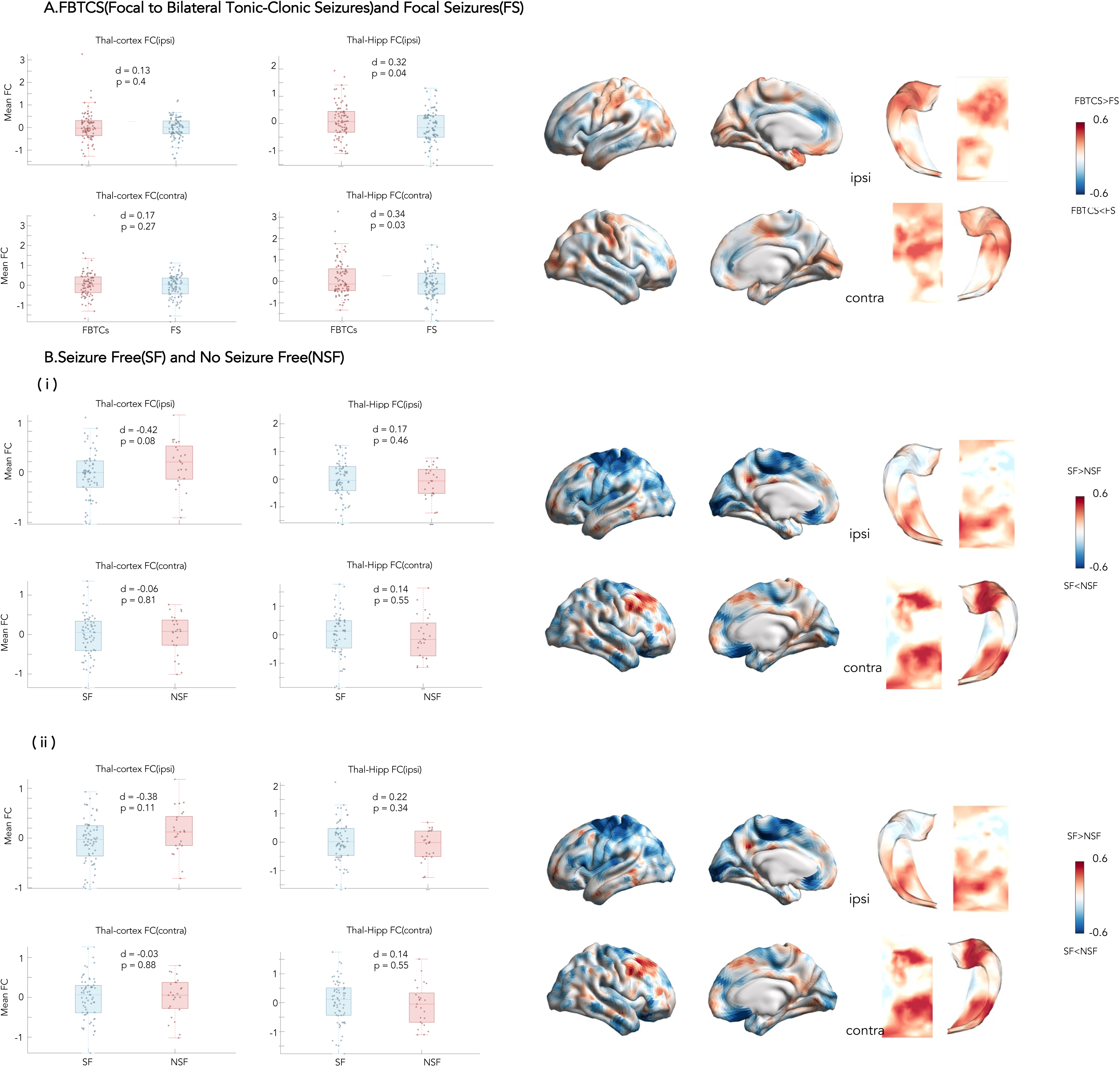

## Notes

### Competing Interest Statement

The authors have declared no competing interest.

