## Supplementary Information for "FUNCTIONAL FRAGMENTATION AND STRUCTURAL DRIVERS OF THALAMO-CORTICAL CIRCUITS IN TEMPORAL LOBE EPILEPSY"

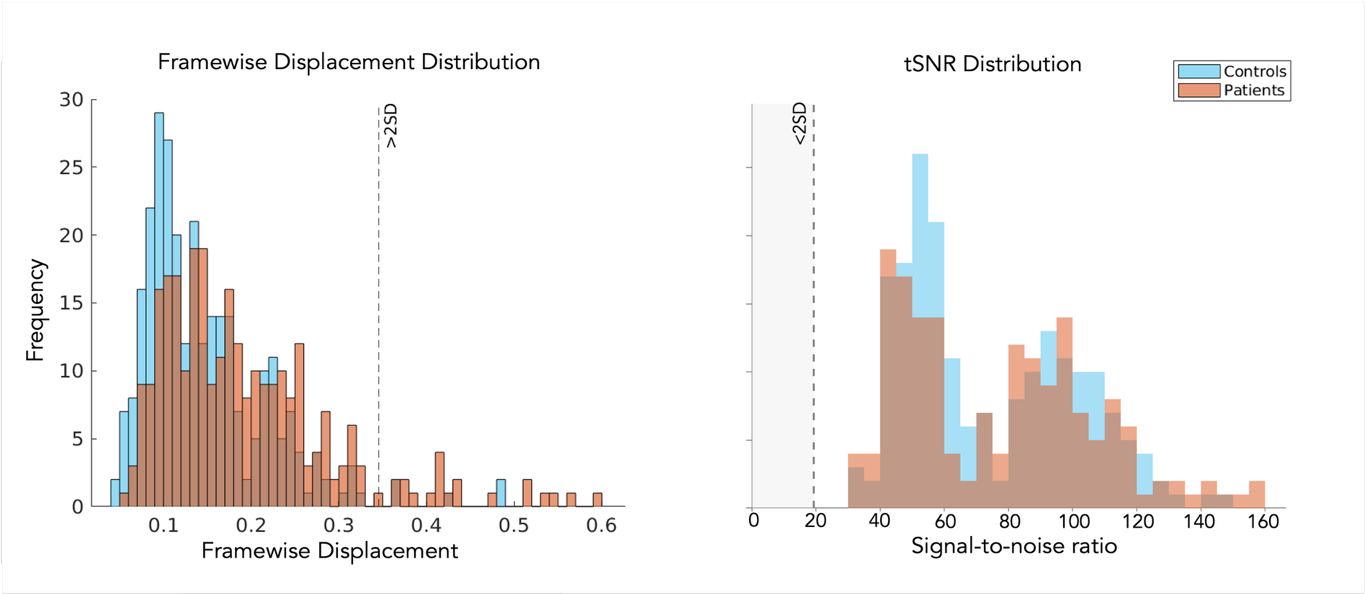

**Fig. S1. Participant inclusion and quality control of the rs-fMRI data.** **(A)** Head motion analysis (participants with motion > 2 SD excluded) **(B)** Signal-to-noise ratio distribution.

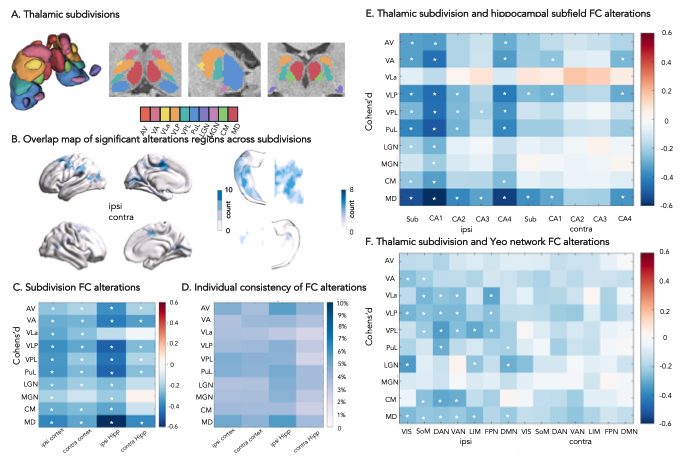

**Fig.S2.** **Validation of subregional FC findings using HIPS-THOMAS atlas.** **(A)** The HIPS-THOMAS atlas segments the thalamus into ten anatomically-defined nuclei. **(B)** Overlap maps of significant thalamo-cortical/hippo campal FC differences (TLE vs HC) across nuclei. **(C)** Generalized linear models, controlling for age and sex, showed FC reductions across multiple nuclei, with MD exhibiting the most pronounced hippocampal FC decrease — consistent with the Tian VAip finding given their high spatial correspondence (**Fig. 2C**). **(D)** Individual-level consistency of FC alterations. **(E)** Nucleus–hippocampal subfield FC alterations, showing the greatest reduction between ipsilateral MD and CA1, replicating the Tian VAip–CA1 pattern (**Fig. 2E**). **(F)** Nucleus–Yeo network FC alterations, with widespread reductions distributed across multiple nuclei (VLP, VPL, CM) and convergent involvement of dorsal attention and default mode networks (**Fig. 2F**). Asterisks denote significance after FDR correction (**p*_FDR_ < 0.05). Abbreviations: AV: anteroventral; VA: ventral anterior; VLa: ventral lateral anterior; VLP: ventral lateral posterior; VPL: ventral posterior lateral; PuL: pulvinar; LGN: lateral geniculate; MGN: medial geniculate; CM: centromedian; MD: mediodorsal.

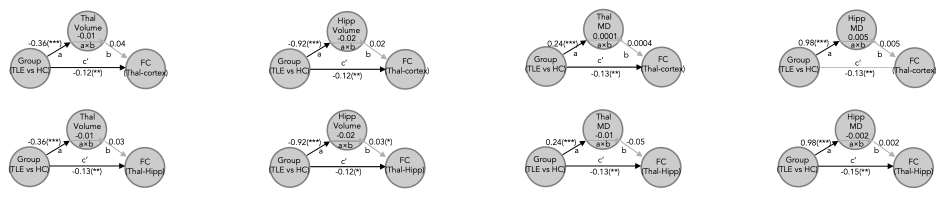

**Fig.S3. Mediation effect on contralateral thalamo-cortical/hippocampal FC.** Mediation analyses using *group* (TLE *v*s HC) as the predictive variable, *structure* (ipsilateral thalamic/hippocampal volume, MD) as the mediator variable, and mean contralateral *FC* in significant alteration regions as the dependent variable. Path *a* indicated the effects of *group* on *structure*, path *b* indicated the effects of *structure* on *FC*, and paths *c*' and *a*×*b* indicated the direct and the indirect/mediation effects of *group* on FC, respectively. *** *p*<0.0005, ** *p*<0.005, * *p*<0.05.

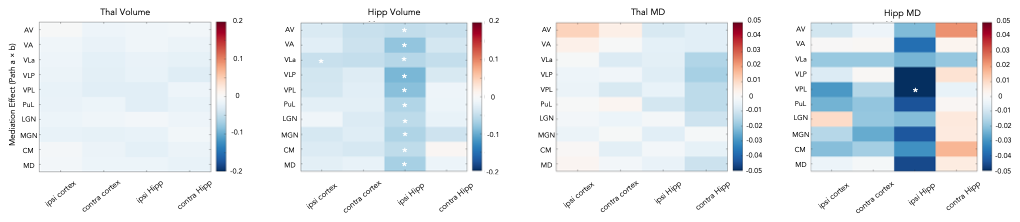

**Fig.S4.** **Validation of subregional FC mediation effects using HIPS-THOMAS atlas.**

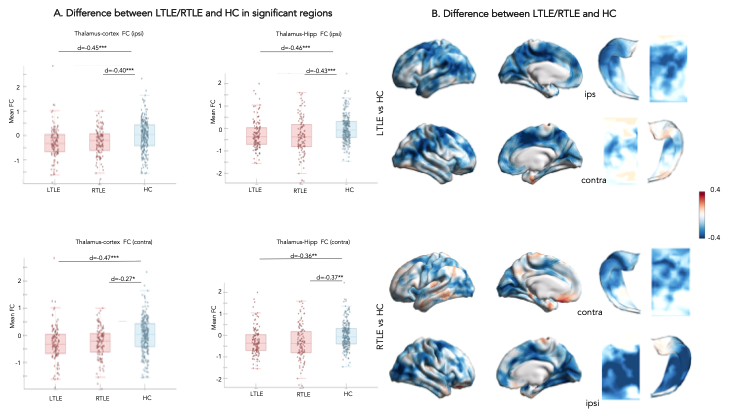

**Fig. S5. Functional connectivity in left and right TLE.** **(A)** Mean FC values comparison between LTLE/ RTLE and HC within significant regions identified in the multi-site analysis. **(B)** vertex-wise comparisons (*** *p*<0.0005, ** *p*<0.005,* *p*<0.05). Both analyses controlled for age, sex, and acquisition sites.

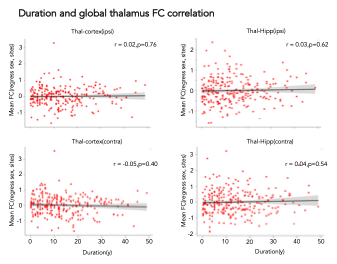

**Fig. S6. The correlation coefficient between FC and duration .** Correlation coefficients between disease duration and global thalamic FC in TLE patients (*n*=250), controlling for sex and acquisition sites as covariates. Abbreviations: *TLE*: temporal lobe epilepsy; *ipsi*: ipsilateral; *contra*: contralateral.

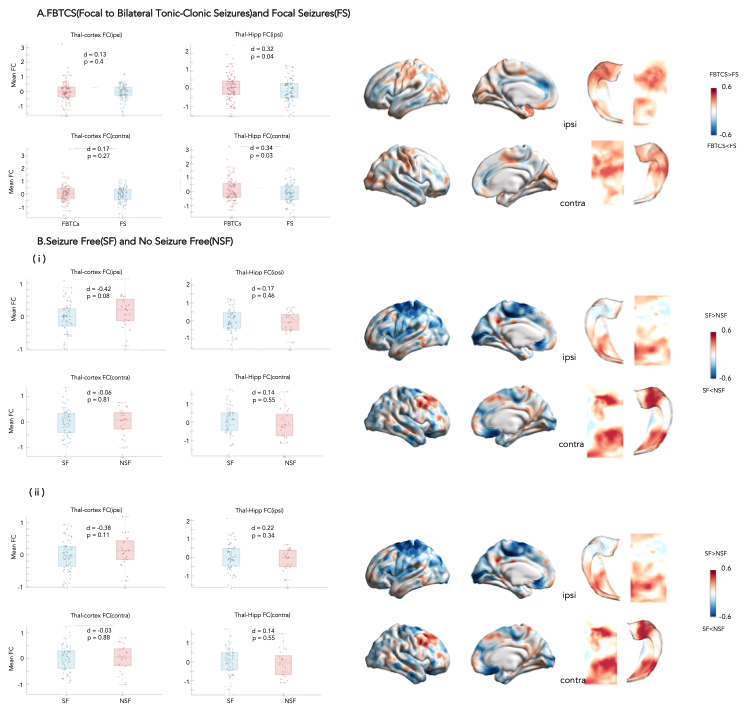

**Fig. S7. Effects of FBTCs and SF/NSF. (A)** Mean FC values in significant regions were compared between patients FBTCs (n = 83) and FS (n = 88) groups, alongside vertex-wise comparisons, with sex and acquisition sites included as covariates in both analyses. **(B)** SF (n=65) vs. NSF (n=25): **(i)** Same analysis as (A); **(ii)** Additional adjustment for disease duration and ipsilateral hippocampal volume. Abbreviations: *TLE*: temporal lobe epilepsy; *ipsi*:ipsilateral; *contra:* contralateral; *FBTCs:* focal to bilateral tonic-clonic seizures; *FS:* focal seizure; *SF:*seizure free; *NSF:*non-seizure free.
